# Metagenomic binning with assembly graph embeddings

**DOI:** 10.1101/2022.02.25.481923

**Authors:** Andre Lamurias, Mantas Sereika, Mads Albertsen, Katja Hose, Thomas Dyhre Nielsen

## Abstract

Despite recent advancements in sequencing technologies and assembly methods, obtaining high-quality microbial genomes from metagenomic samples is still not a trivial task. Current metagenomic binners do not take full advantage of assembly graphs and are not optimized for long-read assemblies. Deep graph learning algorithms have been proposed in other fields to deal with complex graph data structures. The graph structure generated during the assembly process could be integrated with contig features to obtain better bins with deep learning.

We propose GraphMB, which uses graph neural networks to incorporate the assembly graph into the binning process. We test GraphMB on long-read datasets of different complexities, and compare the performance with other binners in terms of the number of High Quality (HQ) genome bins obtained. With our approach, we were able to obtain unique bins on all real datasets, and obtain more bins on most datasets. In particular, we obtained on average 17.5% more HQ bins when compared to state-of-the-art binners and 13.7% when aggregating the results of our binner with the others. These results indicate that a deep learning model can integrate contig-specific and graph-structure information to improve metagenomic binning. GraphMB is available from https://github.com/MicrobialDarkMatter/GraphMB.

## 1 Introduction

Microbial communities play a vital role in most processes in the biosphere and are essential for solving present and future environmental challenges [1]. Examples include the impact of the human microbiome on health and disease [2], discovery of new antibiotics [3], and turning waste products into valuables [4]. Metagenomics holds the promise to enable access to genomes of microbes from complex microbial communities and thereby aid to realize their potential. However, high-quality genomes are difficult to obtain from complex communities, since it is not trivial to determine which DNA sequences originate from the same microbial genome.

To retrieve metagenome assembled genomes (MAGs) from complex metagenomes, several binners have been proposed based on composition and abundance features [5]. Composition refers to the k-mer frequencies of a particular contig and can be used to distinguish between different species [6]. The abundance (coverage) of a contig in one or more samples has also been shown to be a powerful feature to retrieve MAGs [7, 8, 9], which is usually referred to as differential abundance (or coverage) binning.

One of the most successful binners is MetaBAT2 [10]. It uses coverage and composition to compute a pairwise distance matrix for all contig pairs, with the composition feature based on an empirical posterior probability calculated from a set of reference genomes. A graph-based clustering algorithm is then used to bin the contigs based on their distances, where the contigs are linked according to their similarity scores. Wu et al. [11] presented a similar method, MaxBin2, that uses an Expectation-Maximization algorithm to estimate the probability of a contig belonging to a particular bin, but also used single-copy marker genes to estimate the number of bins. Although more composition and abundance methods have been proposed [12, 13, 14, 15, 16], these two can be considered the most established and commonly used.

More recently, deep learning-based methods have been used to improve metagenomic binning. Deep learning models present an advantage over other statistic methods since this type of model can learn complex patterns in the data that would be difficult to manually model with other methods. Nissen et al. [17] proposed, VAMB, a binner based on a variational auto-encoder to encode composition and abundance features into low dimension embeddings that can lead to better binning results on the datasets tested. Other deep learning approaches have also been recently proposed. LRBinner [18] adapts variational auto-encoders to long-reads, while SemiBin [19] uses a semi-supervised siamese neural network with must-link and cannot-link constraints obtained with reference genomes.

Some recent works have also used the assembly graph to improve metagenomic binning. The common assumption is that contigs that were linked on the assembly graph should also be binned together, as they are likely broken into contigs based on internal genome repeats. Mallawaarachchi et al. [20] presented a method that refines bins from other tools using information from the assembly graph. Their method, GraphBin, refines the clusters of contigs that were separated by binning but were linked in the assembly graph. They navigate the assembly graph using a label propagation algorithm to refine the binning. MetaCoAG also uses the assembly graph for post-processing bin refinement [21].

However, both GraphBin and MetaCoAG only use the assembly graph during post-processing, instead of integrating into the binning process. This means that their clustering algorithm uses only contig-specific features, ignoring the connectivity information provided the assembly graph until after an initial clustering is obtained. By integrating the assembly graph only as a post-processing step, more errors can be introduced if this information is not properly used, i.e. contigs may be incorrectly assigned to bins due to misleading links in the assembly graph. This is more likely to occur in complex samples where multiple strains occur and contigs will be connected in the assembly graph even if they belong to different but similar genomes.

With the recent successes of deep neural networks in various problems, there has also been an increasing focus on adapting those approaches for graph data structures. Graph Neural Networks (GNNs) take advantage of the connectivity information in a graph and can be used to perform node, edge and graph-level tasks. The GraphSAGE [22] algorithm samples neighbors from each node and updates the node’s embeddings taking into account the embeddings of its neighbors. The embeddings of the neighbors are aggregated and concatenated with the node embeddings. The resulting vector is the input for the next layer, and the sampling process is repeated. To train GraphSAGE on unlabeled nodes, the similarity between neighboring nodes is calculated and the model weights are updated in order to maximize this similarity, while minimizing the similarity between random pairs of nodes. The loss function used is a binary cross-entropy function, that takes as input the dot product between the embeddings of the two nodes of an edge. However, the random negative sampling strategy is not optimal for assembly graphs, since two disconnected contigs may also belong to the same genome. Furthermore, the original GraphSAGE implementation also considers all neighbors to be of the same importance, while on an assembly graph some edges may be stronger than others.

Finally, most binning methods are developed only on short-read assemblies [5], and only very few binners have been developed with a focus on long-read assemblies [18, 23]. While long-read sequencing technologies have gained traction, there is still a lack of benchmarks and studies on long-read sequencing for metagenomics [24, 25, 26, 27]. The longer read length results in much improved assemblies that generates more sparse assembly graphs and enables more robust estimations of composition and coverage.

Here we present GraphMB, a binner developed using long-read metagenomic data and incorporates the assembly graph into the contig features learning process, taking full advantage of its potential by training a neural network to give more importance to higher coverage edges. The graph-aware features of each contig are based on its own features, as well as on the contig-specific features of its neighboring contigs. We accomplish this using state-of-the-art deep learning techniques, in particular Graph Neural Networks (GNN), a type of deep learning model that can learn representations of graph nodes based on node features and the graph structure.

## 2 Materials and Methods

The main idea behind GraphMB is to generate embeddings based on contig-specific features and the assembly graph, which are then clustered into bins and evaluated according to completeness and contamination. The advantage of clustering embeddings instead of the original features is that these embeddings are of a smaller dimension and can encode relationships that are latent in the original features. We improve upon existing binners by incorporating the assembly graph into the training process. The assembly graph describes which contigs are connected, and how many reads support that connection (read coverage). We use this information to train a GNN, and generate embeddings that take into account the neighborhood of a contig. Figure 1 provides an overview of GraphMB, and the following sections explain each step of the process.

**Figure 1:**
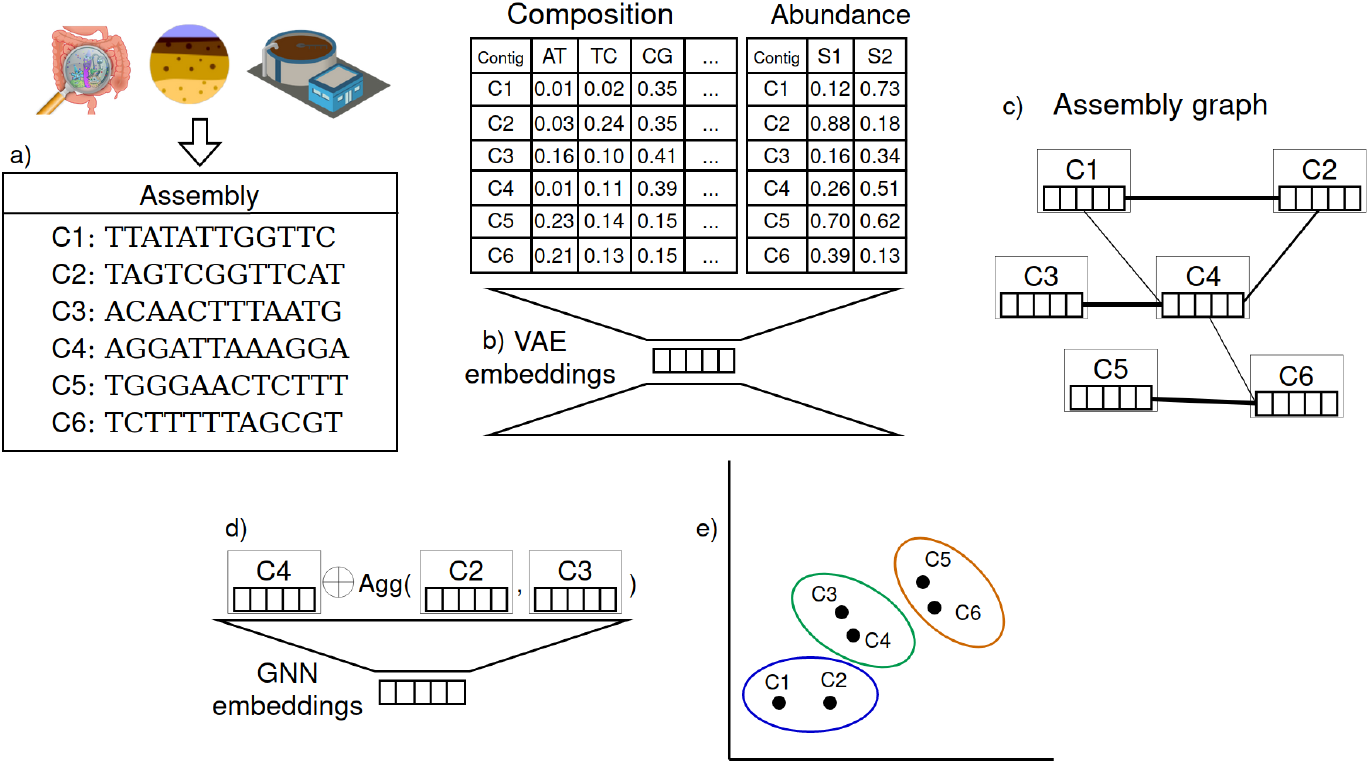
GraphMB’s workflow. (a) The metagenome of an environmental sample is sequenced and assembled into contigs; (b) Initial embeddings are computed with a variational auto-encoder based on k-mer composition and abundance features; (c) The input of the GNN are the initial contig embeddings and the graph structure provided by the assembly graph. The thickness of the edge corresponds to the number of reads that cover it. (d) The GNN model learns new embeddings by aggregating neighboring contigs (nodes in the assembly graph). (e) The final embeddings are clustered and bins are obtained.

### 2.1 Input data

GraphMB requires an assembly consisting of a set of contig sequences in FASTA format and an assembly graph in GFA format. We tested exclusively with assemblies generated with Flye [24] (v2.9), which has the advantage of including the read coverage of each edge into the assembly graph file. Additionally, a Comma-separated Values (CSV) file with the essential single copy marker genes found on each contig may be provided to select the best training epoch. To take advantage of the same abundance features as other binners, another CSV file with the abundance of each contig on different samples may be provided. The format of this file is one row per contig and the mean base coverage and variance on each sample as columns, which is compatible with other binners (MetaBAT2 and VAMB).

### 2.2 Contig-specific embeddings

We first generate contig embeddings using a Variational Auto-Encoder (VAE) model that takes as input composition and abundance features. This model defines a multivariate distribution over a latent representation of the features (Figure 1b). The composition features are calculated on the provided FASTA file, while The abundance features should be pre-calculated and provided as a CSV file previously described. The VAE model has an encoder and a decoder component. The latent representation is learned by training the encoder with a reconstruction loss, comparing the original inputs to the output of the decoder, and with a Kullback–Leibler (KL) divergence loss, which penalizes distributions that diverge from a standard normal distribution. We used the VAMB implementation of VAEs, that separates the reconstruction loss into two components (composition and abundance), which have different weights (10% to composition and 90% to the abundance) [17].

### 2.3 Neighborhood sampling

We have adapted the GraphSAGE sampling algorithm to make better use of the assembly graph information. An assembly graph *G* is constituted by contigs *C* and adjacency matrix *A*. Each contig *c* ∈ *C* has contig-specific feature vector *x*_*c*_ ∈ *X*, obtained in the previous step, and *A*_*i,j*_ = *rc*(*c*_*i*_, *c*_*j*_), where *rc* is the read coverage, if *c*_*i*_ and *c*_*j*_ are connected in the assembly graph, or 0 otherwise. We use the read coverage of each edge as a way to distinguish between pairs of contigs that are more likely to belong to the same genome. If a contig is disconnected from the graph, we pick a random contig as a negative edge. However, if a contig is connected to multiple other contigs, we use the read coverage as a probability of picking an neighbouring edge as a positive edge, and its inverse as the probability of picking it as a negative. For example, in Figure 1c, C4-C3 is more likely to be sampled as a positive edge than C4-C6, since the former has a higher read coverage. This way, the model minimizes the distance between embeddings of pairs of contigs that are connected by high coverage edges.

### 2.4 Graph embeddings

The hidden state of each contig (represented in Figure 1d by the empty squares) is concatenated with the aggregation of the hidden states of the sampled neighbors. Then a feed-forward neural network is trained to generate graph embeddings using the previous concatenation as input. The initial hidden states correspond to the contig-specific features, while for each layer of the network, the hidden states correspond to the output of the previous layer for each contig. The output of the final layer corresponds to the graph embeddings.

We used a loss function that takes advantage of the read coverage information provided by the assembly graph. For the positive edges, we multiply the dot product between the two node embeddings by the normalized read coverage. This way, low coverage edges, which are more likely to introduce noise into the model, will have less impact on the loss function, and we give more importance to the edges with high coverage while training. Therefore, the loss we used is given by:

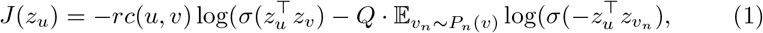

where *z*_*u*_ and *z*_*v*_ are the embeddings of two contigs with *rc* read coverage, and *v*_*n*_ is a randomly sampled negative edge for contig *u*, *P*_*n*_ is the negative sampling distribution previously explained and *Q* the number of negative samples, since multiple negatives can be sampled for each positive edge.

### 2.5 Clustering

We cluster the concatenation of the contig-specific embeddings and graph embeddings with the iterative medoid clustering algorithm used by VAMB, also similar to the one used by MetaBAT2. We cluster the concatenation of both embeddings since we observed that this strategy worked better than clustering only one type of embeddings. This algorithm takes a random seed contig and calculates its embedding distance to all other contigs. Then it uses an iterative process to determine the best medoid contigs and generates clusters with the other contigs that are closest to the medoid. This method has the advantage of not requiring the number of clusters as input, and being easily parallelizable.

### 2.6 Experimental setup

We run experiments on one simulated dataset, six Wastewater Treatment Plant (WWTP) datasets, and one soil sample. As long-read datasets are not part of the benchmarks used by other binners, we simulated our own data. The simulated dataset was generated using badread [28] (v0.2.0), by generating reads according to the methodology proposed by Quince et al. [29]. We simulate reads from 100 strains, corresponding to 50 species, with randomly generated abundances. We then assembled the reads with metaflye v2.9 [24] and ran other binners for comparison. The details of the assembly of each dataset are given on Table 1. The WWTP datasets come from a previous study [26] (Accession number PRJNA629478), from which we used a subset of 6 plants. For each one of the WWTP datasets, we calculated contig coverage on the long-reads used to generated those contigs, as well as three additional short-reads datasets from the same plant but different time points. We assembled with metaflye, and ran three Racon (v1.3.3) polishing rounds with the long-reads and one round with shortreads. Finally, we also tested on a soil sample that originated from a previous study [30] (Accession number PRJEB50688). We developed and optimized the hyperparameters of the network on all datasets, except Damh and Hade which we used to confirm if the model was over-optimized for the other datasets. We have made all datasets available at https://doi.org/10.5281/zenodo.6122610.

**Table 1:** Summary of the datasets used to compare binners. Total size refers to the total number of base pairs in the dataset; Reads N50 is the N50 length of reads; Assembly length refers to the sum of the length of all contigs; Contig N50 is the N50 value for contigs; Mean cov. refers to the mean base coverage of all contigs; Contigs and edges refers to the number of contigs of each assembly and edges in the assembly graph; Samples is the number of samples available to calculate abundance.

| Datasets | Total size (Gbp) | Reads N50 (kbp) | Assembly length (Gbp) | Contigs N50 (kbp) | Mean cov. | Contigs | Edges | Samples |
| --- | --- | --- | --- | --- | --- | --- | --- | --- |
| Strong100 | 7.5 | 13.3 | 0.17 | 175.0 | 42 | 852 | 670 | 1 |
| Hjor | 16.0 | 8.7 | 0.86 | 80.4 | 13 | 19496 | 5937 | 4 |
| Viby | 17.2 | 14.0 | 1.32 | 101.0 | 7 | 23389 | 7800 | 4 |
| Damh | 26.7 | 14.3 | 1.93 | 119.0 | 8 | 32771 | 14066 | 4 |
| Mari | 23.3 | 10.1 | 1.69 | 83.1 | 8 | 36611 | 12651 | 4 |
| AalE | 27.7 | 10.2 | 1.92 | 83.4 | 8 | 40827 | 12425 | 4 |
| Hade | 45.2 | 9.8 | 3.01 | 73.9 | 9 | 70402 | 27952 | 4 |
| Soil | 115.0 | 7.7 | 1.98 | 93.3 | 19 | 51135 | 69522 | 1 |

### 2.7 Evaluation

We compared the results of our binner with five other binners on the same datasets, using their default values. All binners we used take as input the contig sequences and their abundances. We used MetaBAT2 [10], since it had obtained good results on the WWTP datasets, and is generally considered the state-of-the-art on recent papers [31, 32]. We also used MaxBin2 [11], another established metagenomic binner. VAMB [17] is a deep learning-based binner, which we compare with our approach. GraphBin [20] is a binner that also takes advantage of the assembly graph but has only been tested on short-read assemblies. We run Graphbin with the output of MetaBAT as initial bins, which are required by this tool. Finally, we ran SemiBin [19], a recently proposed deep learning binner, using one of the pretrain models provided by the authors (ocean model) as well as training on our own data with the default parameters.

Each bin generated by GraphMB and other binners was evaluated for completeness and contamination with CheckM [33] (–reduced-tree, version 1.1.2) and dereplicated using dRep [34] (version 2.3.2). We considered High Quality (HQ) bins as those with *>*90 completeness and *<*5 contamination. dRep generates bin clusters based on multiple sets of bins obtained with different approaches. The bin clusters contain bins from different approaches that are similar enough to be clustered together. We consider unique bins as those that are HQ and were not clustered together with any other HQ bin from a different approach. Finally, we used DASTool to combine the bins produced by all tools, generating a set of bins that is a combination of all approaches.

## 3 Results

We implemented the proposed method in Python and compared its performance to state-of-the-art binners on simulated and real-world datasets.

### 3.1 Implementation

We implemented GraphMB in Python 3.7, Pytorch 1.10 and DGL 0.6.1. It can run both on CPU (single and multithread) and GPU. The package can be installed from GitHub, using pip, or with anaconda. We provide simple instructions on the GitHub page^1^ including example commands, as well as a link to more detailed documentation. The GitHub page also includes the simulated dataset for testing.

Many parameters can be configured, however, we defined default values for what we used in our experiments. Some parameters, such as the size of the embedding dimension and batch size, can be set automatically according to the size of the input datasets. GraphMB has three graph convolution layers, with hidden dimension of 512 and output dimension of 64, learning rate of 0.00005 and ReLU activation function. We trained each model for 100 epochs.

The output of GraphMB is a TSV file mapping each contig to a bin. The model and embeddings of the last epoch are saved to disk. If a contig marker file is provided, GraphMB also saves the model that obtained the best performance on those markers, which differs from the final CheckM evaluation, but is still a good indication of the best epoch to stop model training, and we used this criterion for the results shown. The training process can also be stopped if the previous two epochs did not reduce the loss by more than a certain threshold. We do not filter by bin size or write the contigs to file by default, since this can be accomplished with a post-processing script, and may not be required for all applications.

### 3.2 Comparison to other binners

Table 2 shows the results obtained for all datasets by all tested binners. GraphBin, MaxBin2, SemiBin-ocean, Semibin-train, VAMB, MetaBAT2, and GraphMB refers to the number of HQ bins obtained with each approach for each dataset.

**Table 2:** Results obtained with GraphMB and state-of-the-art binning tools.. The WWTP datasets are sorted by ascending size of assembly in terms of number of contigs. The Soil dataset is separate because it has a much higher complexity than the WWTP datasets

| HQ bins | Strong100 | Hjor | Viby | Damh | Mari | AalE | Hade | Soil |
| --- | --- | --- | --- | --- | --- | --- | --- | --- |
| GraphBin | 30 | 11 | 15 | 14 | 16 | 12 | 6 | 0 |
| Maxbin2 | 27 | 12 | 19 | 16 | 14 | 12 | 19 | 0 |
| SemiBin-ocean | 30 | 11 | 1 | 22 | 18 | 21 | 7 | 0 |
| SemiBin-train | 27 | 7 | 4 | 23 | 22 | 32 | 25 | 0 |
| VAMB | 28 | 22 | 12 | 22 | 30 | 37 | 28 | 0 |
| MetaBAT2 | 32 | 23 | 29 | 41 | 39 | 43 | 44 | 2 |
| **GraphMB** | 33 | 25 | 23 | 43 | 48 | 46 | 52 | 3 |
| $\Delta$ VAMB | 5 | 3 | 11 | 21 | 18 | 9 | 24 | 3 |
| $\Delta$ MetaBAT | 1 | 2 | -6 | 2 | 9 | 3 | 8 | 1 |
| $\Delta$ % VAMB | 15.2% | 12.0% | 47.8% | 48.8% | 37.5% | 19.6% | 46.2% | 100.0% |
| $\Delta$ % MetaBAT | 3.0% | 8.0% | -26.1% | 4.7% | 18.8% | 6.5% | 15.4% | 33.3% |
| GraphMB dRep unique | 0 | 1 | 2 | 4 | 6 | 8 | 12 | 2 |
| DASTool w/o GraphMB | 37 | 32 | 32 | 41 | 43 | 43 | 51 | 15 |
| DASTool w/ GraphMB | 37 | 33 | 32 | 46 | 47 | 48 | 58 | 16 |

Table 2 also shows the difference in terms of number of HQ bins between GraphMB and both VAMB and MetaBAT2, in absolute value and in percentage. We focus our comparison between GraphMB and those two since MetaBAT2 obtained the best results on most datasets, and VAMB is the closest to our approach. “GraphMB dRep unique” refers to how many of the HQ bins generated by GraphMB were not matched with HQ bins from the other binners according to the dRep analysis. dRep finds bins from different binning approaches that correspond to essentially identical genomes. The number on the table corresponds to groups of bins that have only one HQ bin, and that bin was obtained with GraphMB, i.e. HQ bins that only GraphMB could identify.

We obtained more bins using our graph embedding method when compared to VAMB. For the other WWTP datasets, we obtained between 3 and 21 more HQ bins (12%-49%) on the WWTP datasets, in comparison to VAMB. Compared with MetaBAT2, we obtained between 1 and 9 more HQ bins (3%-22%), and our approach obtained more bins on all but one of the real datasets. It did not outperform MetaBAT2 on one of the WWTP datasets, where VAMB also obtained lower results. The GraphBin approach obtained worse results than the other binners, indicating that this particular graph-based approach is not optimized for the long-read assemblies of these datasets. We observed that GraphBin incorrectly merged many bins, obtaining highly contaminated bins. The SemiBin-ocean approach, also obtained a low number of HQ bins on most datasets, possibly since the pretrained model used was also trained on short-read assemblies. However, while training SemiBin on our own data (one model for each dataset), we found that it only improved the results in some cases, indicating that additional hyperparameter tuning may be necessary.

### 3.3 Ensemble results

After combining multiple binning results with dRep, we observed that our approach was able to recover HQ bins that were not recovered by other approaches. This corresponded to a total of 35 bins across all datasets. Our approach obtained distinct bins from other others on all the real-world datasets.

We used DASTool [35] to observe if our approach could improve the aggregation of bins obtained from multiple approaches. We first combined the output bins of MetaBAT2, MaxBin2, GraphBin and VAMB, in order to include a variety of approaches, and then the same bins but also the output bins of GraphMB. This resulted in more HQ bins than any of the individual binners in most cases. Since we obtained unique bins on all datasets, we expected that combining our method to others would also results in more bins, which was the case for 6 out of 7 real datasets we tested on (6 WWTP datasets plus soil dataset). Using DASTool to aggregate the bins of GraphMB with the others resulted in 23 more HQ bins to be recovered. Furthermore, in four datasets, the aggregation of the other bin sets still obtained fewer HQ bins than GraphMB. Note that the difference between the number of bins obtained with DASTool including GraphMB and excluding it is not the same as the dRep unique GraphMB bins. While DASTool aggregates bins from different approaches, improving theirs scores, dRep only matches the outputs of different approaches, without attempting generate a new bin.

### 3.4 Computational performance

We tested GraphMB both on CPU and GPU environments. We did not account for the assembly and abundance calculation times, since these are preprocessing steps common to all approaches. For small datasets such as the simulated dataset we used, GraphMB can run on CPU, single or multi-threaded. On a single thread, the simulated dataset took about 4 minutes to process. For bigger datasets, we recommend using a GPU. We observed that the processing time scales linearly with the number of contigs, as the Hjor dataset, which is the smallest of the real datasets, took 52 seconds per epoch, and the Hade dataset, the biggest one, took 160 seconds per epoch (see Table 1 to compare graph sizes). We run our experiments on a single Tesla V100 GPU with 32GB RAM. The batch size parameter can be adjusted if less memory is available.

## 4 Discussion

This paper presented GraphMB, a metagenomic binner developed for long-read assemblies, that takes advantage of the assembly graph generated during the assembly process to obtain neighborhood-aware embeddings. These embeddings are used to bin contigs and obtain HQ MAGs. We demonstrated our approach on both simulated and real datasets of diverse complexities. While on the simulated dataset GraphMB worked on par with the other binners, it was able to obtain more HQ bins on the real datasets. Furthermore, it also obtained unique bins that other binners could not recover.

The performance of GraphMB depends on the assembly graph, which we can observe when comparing the different datasets we used. We can see that on the soil dataset, which has an assembly graph with more edges, GraphMB obtained lower results, even if higher than the other binners. We intend to adapt Graph Attention Networks [36] to deal with more complex graphs. This type of algorithm learns an attention mechanism to decide which neighbors of a node should have more weight when computing its embedding. This attention mechanism could also be combined with the edge coverage information that we make use of on GraphMB.

GraphMB is also dependent on the quality of the contig-specific embeddings, since these are used as input features to the Graph Neural Network. For example, GraphMB performed worse in comparison to MetaBAT2 in the Viby dataset, where VAMB, which uses only contig-specific embeddings, also had relatively bad performance. To overcome this issue, we plan to implement an end-to-end architecture where the variational auto-encoder could be trained at the same time as the GNN This would mean that instead of having static contig-specific embeddings, these could be fine-tuned while training the GNN.

## Acknowledgements

We would like to acknowledge Caitlin M Singleton for helping with the WWTP raw datasets.

## Footnotes

1 https://github.com/MicrobialDarkMatter/GraphMB

