## Supplementary Material for "Metagenomic binning with assembly graph embeddings"

February 25, 2022

### 1 Strong100 dataset description.

The strong100 dataset is a simulated dataset of long-read with a total of 100 strains from 45 species, where 20 species were represented by a single strain, and the others by multiple strains. We used the same species from [1] and the same genomes whenever possible. The full list of genomes used is provided as supplementary material. We downloaded those genomes and generated random coverages $y_s$ for each species $s$ based on a log-normal distribution, and normalized so that the sum of all $y_s$ was 1. The number of bases simulated for each genome $g$ belonging to species $s$ was given by

$$bp_{s,g} = y_s \cdot p_g \cdot N$$

where $p_g$ is sampled from a Dirichlet distribution according to the number of strains and $N$ is the total number of bases to simulated, which we set to 7.5Gbp. We then use badread[1] (v0.2.0) to simulate long-reads of each genome. We set the mean read length to 10000 and standard deviation to 7000, mean, max and stdev identity to 98, 99.9 and 5, respectively, and the error model to the default `nanopore2020 model`. The simulated reads were then assembled with flye in the same way as the reads from real datasets. We provide the code to generate random reads at `https://github.com/AndreLamurias/binning_workflows`.

### 2 Real-world datasets

Table 1 shows the accession number and reference of the datasets used in this study.

### 3 Medium-Quality bins

Table 2 shows the number of MQ bins (>50 completeness and <10 contamination) obtained with each approach, and in comparison to GraphMB.

[1] `https://github.com/rrwick/Badread`

Table 1: Real-world datasets used in this study.

| Name | Accession No. | Reference |
|---|---|---|
| Hjor | SRX8234968, SRX8234918, SRX8234919, SRX8234920 | [2] |
| Viby | SRX8234979, SRX8234951, SRX8234952, SRX8234953 | [2] |
| Damh | SRX8234959, SRX8234889, SRX8234890, SRX8234891 | [2] |
| Mari | SRX8234971, SRX8234928, SRX8234929, SRX8234930 | [2] |
| AalE | SRX8234954, SRX8234897, SRX8234912, SRX8234913 | [2] |
| Hade | SRX8234965, SRX8234909, SRX8234910, SRX8234911 | [2] |
| Soil | PRJEB50688 | [3] |

Table 2: Medium-quality (MQ) bins obtained with GraphMB and state-of-the-art binning tools.

| HQ bins | Strong100 | Hjor | Viby | Damh | Mari | AalE | Hade | Soil |
|---|---|---|---|---|---|---|---|---|
| GraphBin | 31 | 78 | 93 | 158 | 110 | 123 | 158 | 63 |
| Maxbin2 | 30 | 41 |  | 75 | 61 | 49 | 76 | 17 |
| SemiBin-ocean | 35 | 53 | 52 | 84 | 84 | 85 | 89 | 5 |
| SemiBin-train | 35 | 38 | 58 | 94 | 91 | 88 | 99 | 0 |
| VAMB | 31 | 79 | 77 | 131 | 126 | 151 | 152 | 2 |
| MetaBAT2 | 38 | 96 | 110 | 204 | 144 | 167 | 206 | 63 |
| **GraphMB** | 33 | 75 | 109 | 191 | 153 | 181 | 232 | 51 |
| Δ VAMB | 2 | -4 | 32 | 60 | 27 | 30 | 80 | 51 |
| Δ MetaBAT | -5 | -21 | -1 | -13 | 9 | 14 | 26 | -12 |
| Δ % VAMB | 1.1% | -5.3% | 29.4% | 31.4% | 17.6% | 16.6% | 34.5% | 100.0% |
| Δ % MetaBAT | -2.8% | -28.0% | -0.9% | -6.8% | 5.9% | 7.7% | 11.2% | -23.5% |
